# Unpacking conditional neutrality: genomic signatures of selection on conditionally beneficial and conditionally deleterious mutations

**DOI:** 10.1101/549329

**Authors:** Jonathan A. Mee, Samuel Yeaman

**Affiliations:** Department of Biology, Mount Royal University, Calgary, Canada; Department of Biological Sciences, University of Calgary, Calgary, Canada

## Abstract

It is common to look for signatures of local adaptation in genomes by identifying loci with extreme levels of allele frequency divergence among populations. This approach to finding genes associated with local adaptation often assumes antagonistic pleiotropy, wherein alternative alleles are strongly favoured in alternative environments. Conditional neutrality has been proposed as an alternative to antagonistic pleiotropy, but conditionally neutral polymorphisms are transient and it is unclear how much outlier signal would be maintained under different forms of conditional neutrality. Here, we use individual-based simulations and a simple analytical heuristic to show that a pattern that mimics local adaptation at the phenotypic level, where each genotype has the highest fitness in its home environment, can be produced by the accumulation of mutations that are neutral in their home environment and deleterious in non-local environments. Because conditionally deleterious mutations likely arise at a rate many times higher than conditionally beneficial mutations, they can have a significant cumulative effect on fitness even when individual effect sizes are small. We show that conditionally deleterious mutations driving non-local maladaptation may be undetectable by even the most powerful genome scans, as differences in allele frequency between populations are typically small. We also explore the evolutionary effects of conditionally-beneficial mutations and find that they can maintain significant signals of local adaptation, and they would be more readily detectable than conditionally deleterious mutations using conventional genome scan approaches. We discuss implications for interpreting outcomes of transplant experiments and genome scans that are used to study the genetic basis of local adaptation.

## Introduction

Local adaptation is the presence of genotype-by-environment interactions for fitness, such that each local population has its highest fitness in its home site (Kawecki and Ebert 2004; Blanquart et al. 2013; Richardson et al. 2014). Whereas reciprocal transplant experiments have revealed extensive evidence for local adaptation at the phenotypic level (Linhart and Grant 1996; Hereford 2009; Blanquart et al. 2013), and numerous genome scans over recent years point to many potential candidates driving local adaptation, there are comparatively few well-studied examples of the genomic basis of local adaptation (reviewed in Song and Mitchell-Olds 2011; Savolainen et al. 2013; Hoban et al. 2016). Most theoretical study of local adaptation has focused on dynamics under antagonistic pleiotropy (AP; Felsenstein 1976; Bürger 2014), whereby the two alleles at a single locus have opposite fitness profiles, with each having high fitness in one environment and low fitness in the other (Figure 1). Under this type of fitness profile, local adaptation tends to be deterministically maintained when the migration rate between the two environments is sufficiently restricted relative to the strength of divergent selection, which yields large and persistent differences in allele frequency between populations (Felsenstein 1976; Burger 2014). But, detailed analyses of common garden experiments have suggested that local adaptation may not depend on antagonistic pleiotropy, and that mutations with a fitness effect in one environment may be neutral in another environment (*e.g*. Fry 1992; Hall et al. 2010; Oakley et al. 2014), termed ‘conditional neutrality’ (Fry 1996; Kawecki 1997; reviewed in Anderson et al. 2013). However, care should be taken in interpreting the lack of a measurable fitness effect for an allele in one environment as conditional neutrality (see discussion, below). Despite the potential role of conditional neutrality in genotype-by-environment interactions for fitness, it is unclear how much signal can be contributed by such mutations at equilibrium, and whether the causal mutations can be easily differentiated from unconditionally neutral loci (see, for example, Yoder and Tiffin, 2018).

**Figure 1.**
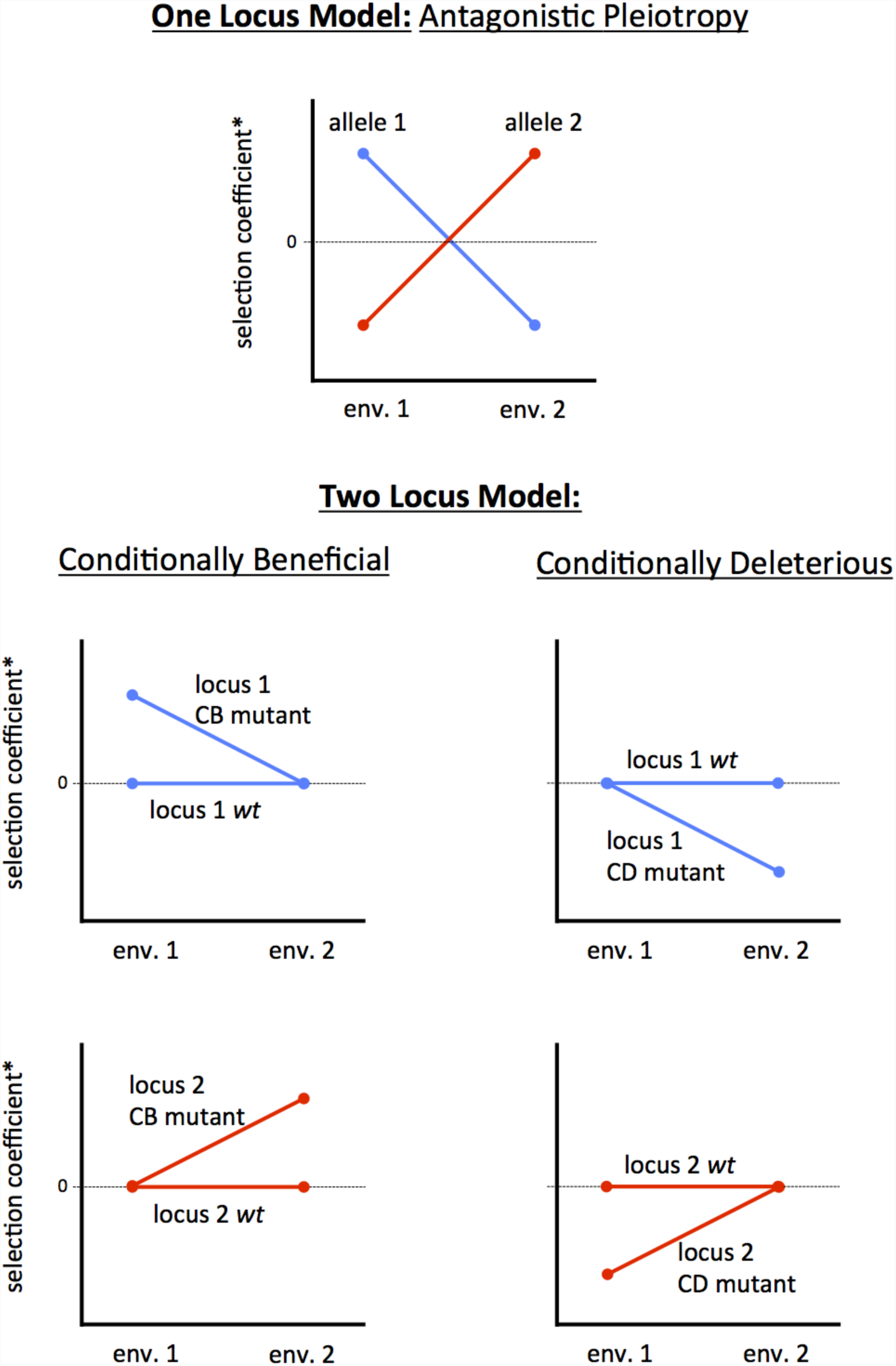
Genotype-by-environment interactions for fitness can arise from several different fitness profiles. The one locus model involves antagonistic pleiotropy. The two locus model (a.k.a. multi-locus model) could involve conditionally beneficial or conditionally deleterious mutations at several loci that collectively result in a phenotypic pattern that is indistinguishable from antagonistic pleiotropy.

There are two fitness profiles involving conditional neutrality that can cause genotype-by-environment interactions for fitness: 1) when new mutations are beneficial in the home site and neutral at other sites (hereafter: “conditionally beneficial” or “CB”), 2) when new mutations are neutral at the home site and deleterious at other sites (hereafter: “conditionally deleterious” or “CD”; Figure 1). There are several important differences between antagonistic pleiotropy and these fitness functions involving conditionally neutral mutations. For antagonistic pleiotropy, polymorphism and local adaptation can be maintained at a deterministic equilibrium for a wide range of migration rates (Felsenstein 1976; Bürger 2014). By contrast, for the CB and CD cases, only monomorphic single-locus equilibria will be stable under deterministic models with high migration; the mutant will replace the ancestral allele in the CB scenario, whereas the mutant will be lost in the CD scenario (Whitlock and Gomulkiewicz 2005; Bürger 2014). Therefore, with sufficient gene flow, polymorphism at any given locus will be transient, persisting only for a short time until the monomorphic equilibrium is regained. In contrast to antagonistic pleiotropy, in order for CB and CD to yield a phenotypic signature resembling local adaptation in a transplant experiment, it is necessary for polymorphisms to segregate at multiple loci with fitness effects conditional in opposite environments (as shown in Figure 1), and new polymorphisms must be renewed continually by mutation. Thus, for both CB and CD, the maintenance of genotype-by-environment interactions for fitness depends upon sufficient spatial structure to delay the eventual return to monomorphism while new CB or CD mutations arise at other loci. Finally, if CD mutations are important drivers of genotype-by-environment interactions, interpreting evidence from common garden or transplant experiments as local adaptation would be misleading because no beneficial mutations are involved. In such cases, CD mutations should more accurately been seen as causing “non-local maladaptation” due to the accumulation of conditionally deleterious mutation load (which we refer to as “non-local load” for simplicity). If we are interested in studying local adaptation because it gives some insight into longer term patterns of species-wide adaptation that drive macroevolution, then it is important to understand the relative contributions of CB, CD, and AP.

A number of recent studies have tested whether observed patterns of allelic differentiation are consistent with conditional neutrality, without differentiating CB and CD mutations (*e.g*. Hall et al. 2010; Fournier-Level et al. 2011; Anderson et al. 2013), likely because distinguishing between these types of mutation in an empirical setting requires identifying which allele was ancestral. Indeed, if only the relative fitness effects of two alleles are considered without any knowledge of which allele was ancestral, it is not possible to discern whether a CD or CB fitness profile is operating. For example, if genotypes *AA* and *aa* have realized fecundities (e.g. number of seeds) of *w*_*AA*,1_ = 20 and *w*_*aa*,1_ = 20 in environment 1 and *w*_*AA*,2_ = 40 and *w*_*aa*,2_ = 50 seeds in environment 2, it is unclear whether *A* is conditionally beneficial or *a* is conditionally deleterious, without knowing which was ancestral. However, the distinction between CB and CD mutations becomes important when considering that universally deleterious mutations occur much more frequently than universally beneficial mutations (Bataillon 2000; Keightley and Eyre-Walker 2010) and that this difference would presumably also extend to their conditionally-dependent counterparts (*i.e*. to CB or CD mutations). Hence, even mutations with very small conditionally deleterious effects (*e.g*. as in the infinitesimal model, *sensu* Bulmer 1980) may accumulate to a sufficient extent to cause substantial levels of non-local load - this appears to be the case for mutational load involving non-conditional mutations (Agrawal and Whitlock 2012).

Evolution by CB mutations should result in a continual increase in absolute fitness at the species scale as CB mutations accumulate and fix. In contrast, patterns of sequence evolution with CD mutations would be dynamically more similar to purifying selection, as the new mutations would eventually be purged such that there would be no gradual increase in absolute fitness. If there are high mutation rates, small effect sizes, and short sojourn times for CD mutations, there may be very little measurable allele frequency divergence between populations evolving in response to different selective regimes. Hence, CD mutations may be important in explaining cases where no significant allelic differentiation is detected (*e.g*. in a genome scan) despite a significant genotype-by-environment interaction for fitness or variation in fitness-related traits.

In this study, we used individual-based population genetic simulations to study the accumulation of conditionally neutral mutations in two patches connected by relatively high migration (*i.e*. such that the effect of migration exceeds the effect of drift: *m* >> 1/N_e_). While it is also biologically interesting to explore the dynamics of CB and CD mutations when drift is strong relative to migration (i.e. *m* >> 1/N_e_), we expect such mutations to accumulate at a rate that is approximately clock-like in the patch where they are neutral (as per the molecular clock; Kimura 1968). As such, we consider this case only briefly, and focus instead on the somewhat less easily predictable question of whether CB and CD mutations can accumulate when migration is high. To this end, we simulated separate cases with either CD or CB mutations under a range of mean selection coefficients and mutation rates, compared their capacity to generate significant genotype-by-environment interactions for fitness, and examined whether genomic signatures of differentiation (*i.e*. F_ST_) could be used to identify causal loci. To aid the interpretation of our simulation results, we compared the observed level of genotype-by-environment interactions for fitness to simple analytical predictions under simplifying assumptions about evolutionary dynamics.

## Methods

### Analytical prediction for non-local load via CD mutations

A simple approximation for the amount of non-local load contributed by CD mutations at equilibrium can be derived by combining several results from classical population genetics, and assuming a large population size (*i.e.* ignoring drift). Consider a given type of locus (CD_2_) where both alleles (*a* and *A*) have equal fitness in patch 1, allele *A* is deleterious in patch 2, and the rate of mutation from wild type *a* to *A* is µ (with the opposite scenario for locus type CD_1_, where *a* is wild type, and *A* is deleterious in patch 1). Under this scenario, the total non-local load for the population inhabiting patch 1 (which would be realized if its inhabitants migrated to patch 2), can be calculated as the summation across all *n* CD_2_ loci:

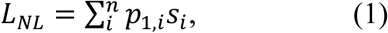

where *s*_*i*_ is the selection coefficient and *p*_*i*,1_ is the frequency of *A* at the *i*^th^ locus in patch 1 (and *q*_*i*,1_ = frequency of *a* in patch 1), with a similar formula applying for patch 2 by summing across CD_1_ loci.

To estimate the equilibrium frequency of CD_2_ mutations in patch 1 (*p*_*i*,1_), we can assume that new mutations that occur in patch 2 will contribute comparatively little to the evolutionary dynamics among the two patches (because they will be purged from patch 2 quickly after arising) and can be ignored. Mutations from *a* to *A* that occur in patch 1 will behave neutrally within the patch, but patch 1 will experience persistent immigration of alleles from patch 2 at rate *m*, all of which will be *a* because of the above assumption that *q*_*i*,2_ = 1 (*i.e.* we are ignoring the short periods of time when a recently mutated *A* in patch 2 has not yet been purged). Because migration will exert a deterministic forcing of allele frequencies in much the same way as natural selection in classical single-population models (*i.e*. the frequency in the next generation, *p*’ = *p*(1 - *m*)), we can substitute *m* for *s* in Haldane’s classic model of mutation load (*p* ≈ µ/*s*; Bürger 2000), so that *A* will reach a mutation-selection equilibrium at *p*_*i*,1_ ≈ µ/*m*. Substituting this into Eq. (1), then the total non-local load will be:

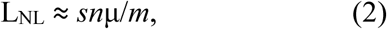

where *s* is the mean selection coefficient.

This is an interesting result, as unlike the classic single-population genetic load that is independent of the selection coefficient where L = *n*µ (Bürger 2000), it predicts that non-local load should depend linearly on *s*. However, this result will tend to underestimate the amount of non-local load because we ignored mutations from *a* to *A* in patch 2, which could migrate to patch 1 and establish before they are purged from patch 2, thereby increasing the non-local load above that accounted for by Eq. (2). Also, some segregation of *A* in patch 2 may occur due to immigration of *A* from patch 1, which will reduce the effect of migration from patch 2 to patch 1 on *p*1 (as some migrants would be *A*), thereby increasing the equilibrium frequency over µ/*m*. This simple approximation may underestimate the non-local load at equilibrium, but provides some insight into how the processes governing non-local load differ from simple genetic load due to unconditionally deleterious mutations.

### Individual-based simulations

To simulate the evolution of individuals distributed in two patches linked by migration, we used SLiM v.1.8 (Messer 2013) with a modification that allowed the inclusion of conditionally neutral mutations (modified code available at https://github.com/samyeaman/slim_condneut). Each individual in these simulations was hermaphroditic and had a diploid genome consisting of ten 100kb chromosomes, for a total of 2,000,000 potential mutational targets (*i.e.* loci) in each genome. The recombination rate was set to 10^−6^ between adjacent loci, except where otherwise specified. At every locus, the ancestral allele was neutral in both patches. When a mutation arose at a given locus, it had a 1/3 chance of being universally neutral, a 1/3 chance of being neutral in patch 1 and selected in patch 2, and a 1/3 chance of being selected in patch 1 and neutral in patch 2. The selection coefficients were drawn from an exponential distribution, with a mean value of *s* specified for each parameter set. In simulations with CD mutations, we specified negative values for mean *s* (*i.e*. multiplying the exponentially-distributed values by −1), and in simulations with CB mutations, we specified a positive value for mean *s*. All mutations were codominant (*h* = 0.5), and mutation effects were additive. It would be interesting to investigate other forms of epistasis (i.e. positive or negative), but such an investigation is outside the scope of this study. The per-locus mutation rate (µ) was set to 10^−8^ or 10^−9^ for simulations involving conditionally deleterious mutations, and to 10^−10^ or 10^−11^ for simulations involving conditionally beneficial mutations. Empirical estimates of deleterious mutation rates are highly variable, but, in humans, the rate is likely bracketed by the mutations rates used in our simulations (Agrawal and Whitlock 2012). The beneficial mutations rates used in our simulations are simply based on an assumption that the rate is at least one or two orders of magnitude lower than the deleterious rate.

The number of individuals per patch (*N*) was set to 1000 or 10,000. The migration rate (*m*) between the two patches was set to 0.5 for the first 50,000 generations of the simulation and then reduced to 0.01 or 0.001 for the last 50,000 generations. The genome of every individual in both patches, including mutational information for every basepair position, was sampled every 10,000 generations. We ran at least 25 replicates of each parameter combination – in a few cases where mutation rates were lowest (10^−10^ and 10^− 11^), we ran additional replicates (up to 100) when there were insufficient mutations across the smaller number of replicates to compute summary statistics.

## Results

### Maintenance of GxE for fitness with conditionally neutral alleles

To compare the extent of genotype-by-environment fitness interactions evolving due to either CD or CB mutations (*i.e*. non-local load in the case of CD mutations, or transient local adaptation in the case of CB mutations), we calculated the average home-away fitness difference over the last 40,000 generations of our simulations. We note that in our two-deme two-habitat model, with symmetric fitness effects, there is little to no difference between “home-away” and “local-foreign” comparisons on average (*censu* Kawecki and Ebert 2004), and both criteria yield similar quantification of local adaptation (supplementary material, Figure S1). In the simulations with CD mutations, home and away fitness tended to stabilize shortly after the reduction in migration rate and before the last 40,000 generations (supplementary material, Figure S2 and S3). Whereas many of the simulations with CB mutations also stabilized before the last 40,000 generations (supplementary material, Figure S4 and S5), fitness tended to increase indefinitely for some simulations with higher values of *s*, and fluctuate extensively for smaller values of *N*, rather than reach an apparent stable state. Hence, our simulation results regarding the dynamics of CD mutations are more easily interpretable than those regarding the dynamics of CB mutations.

To compare the analytical prediction for non-local load from Eq. (2) with results from simulations, we used our calculation of home-away fitness difference as a measure of non-local load. We found that this quantity increased linearly with *n*, *s*, and µ, and decreased linearly with *m*, showing broad agreement with the prediction of Eq. (2) (Figure 2). We note that while the simulations conform to this linear relationship for intermediate strengths of selection, they depart from it for *Nm* = 1 at very low and very high strengths of selection, likely because *Nm* = 1 approaches the region of parameter space where drift dominates over migration, causing the two patches to evolve independently and mutations to accumulate at more of a clock-like rate in the patch where they are neutral (*i.e*. with much delayed purging). These patterns were unaffected when we varied the number of mutable sites, the number of chromosomes, or the recombination rate in our simulations (supplementary material, Figure S6 and S7).

**Figure 2.**
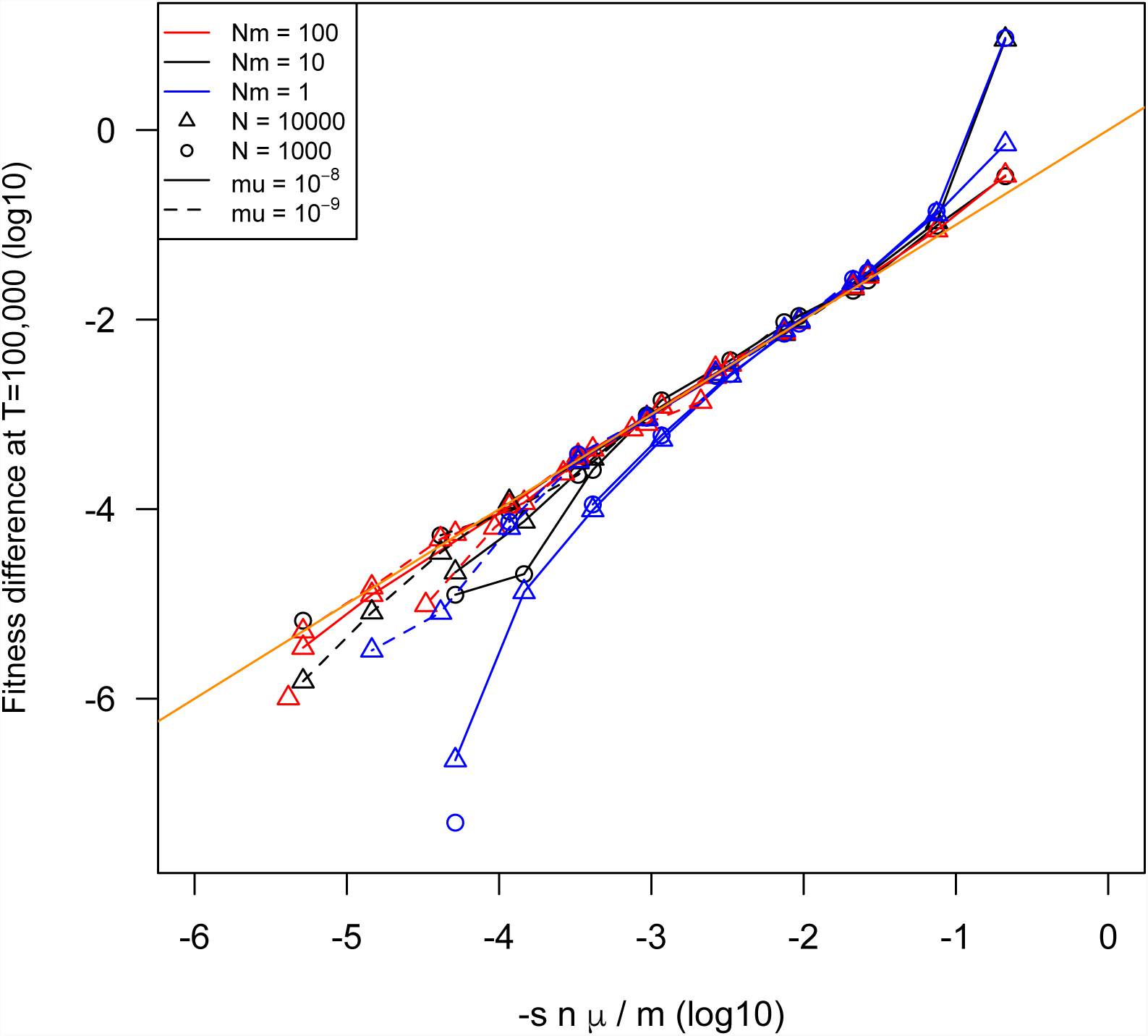
Non-local load driven by CD mutations after 100,000 generations, compared to the prediction from Eq. (2). Orange solid line shows the y = x line.

Whereas there was no apparent or predicted effect of population size on home-away fitness differences in the case of CD mutations (Figure 2 & 3), home-away fitness differences for CB mutations increased noticeably from N = 1000 to N = 10,000 for equivalent values of *m*, µ, and *s* (Figure 3). This likely occurred because the rate of accumulation of positively selected mutations in a single population is approximately 4*Nn*µ*hs*, based on Haldane’s classic result for fixation probability of 2*s* (Haldane 1927) and a mutation rate of 2N*n*µ, which scales linearly with *N*. This suggests that the transient beneficial fitness effect should also scale with *N* as long as the rate of fixation in the patch where alleles are neutral is independent of *N*. This seems likely because once the allele is established where it is beneficial, migration will have a deterministic forcing effect on the allele frequency where it is neutral. It would be interesting to derive analytical results to explore this dynamic, but this is beyond the scope of this paper.

**Figure 3.**
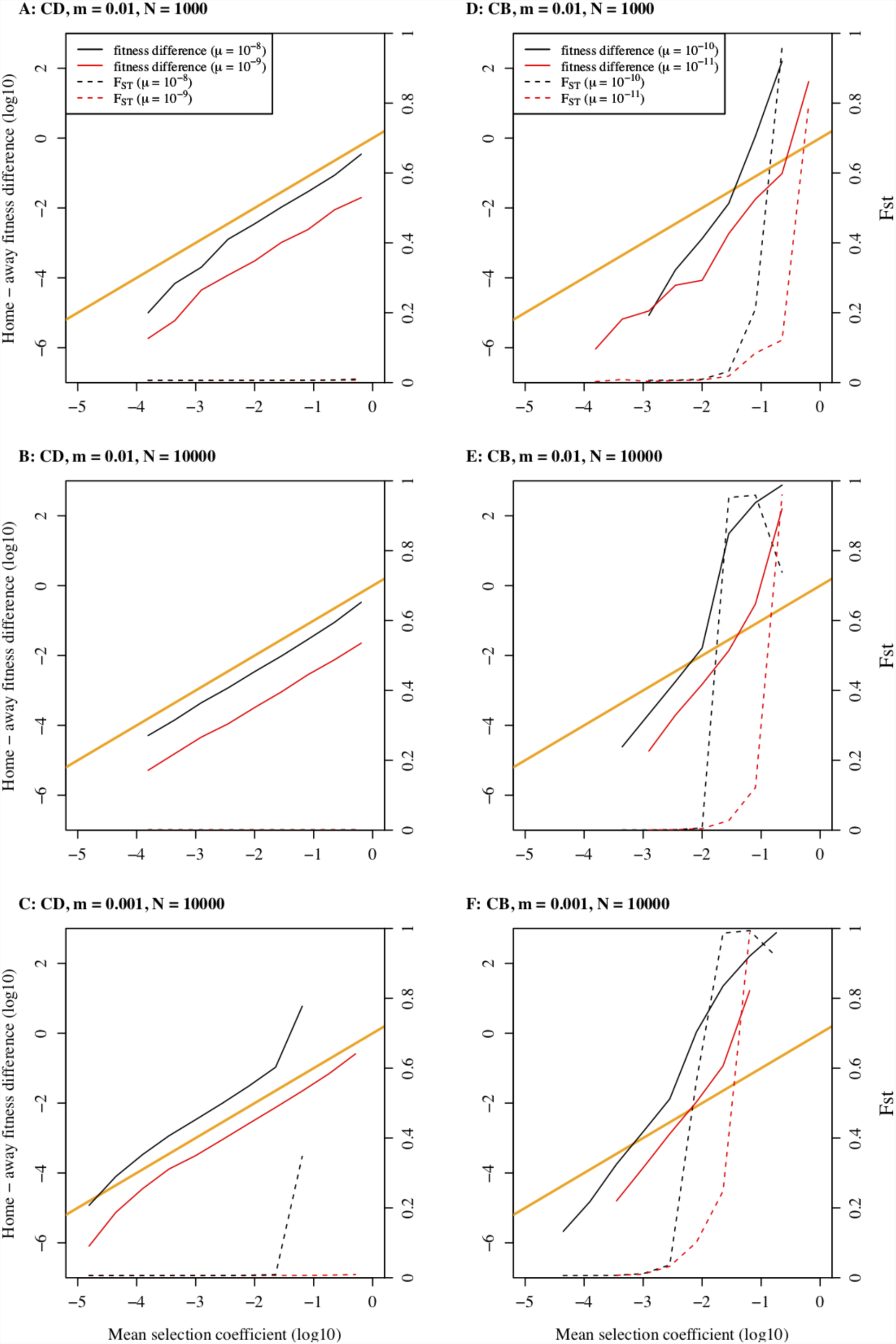
The individual effects of migration rate (*m*), population size (*N*), mean selection coefficient (*s*), and mutation rate (µ) on non-local load driven by CD mutations (A-C) differed from their effects on transient local adaptation driven by CB mutations (D-F) in our simulations. The orange line is the y = x line for the relationship between fitness difference and selection coefficient. The dashed lines show mean allelic differentiation between patches across all loci (F_ST_, right-hand axis) for each parameter combination. Fitness differences were averaged over the last 40,000 generations, and it should be noted that while simulations with CD mutations had reached a stable state where this value was not changing over time, many simulations with CB mutations were not at equilibrium, but were continuously accumulating new mutations with gradually increasing fitness differences.

To compare overall genomic divergence resulting from CD and CB mutations, we also calculated the average allele frequency differentiation between patches across all loci (F_ST_) over the last 40,000 generations of our simulations. The average allele frequency differentiation between patches increased to a greater extent with CB mutations relative to CD mutations (Figure 3). In most cases, non-local load accumulated in the absence of any detectable increase in F_ST_ (Figure 3 A-C). In contrast, there was very little parameter space in the simulations with CB mutations where local adaptation occurred without an increase in F_ST_ (Figure 3 D-F).

### Non-local maladaptation when migration is weak relative to drift

While we mainly focused on cases where migration was strong relative to drift (*i.e*. *m* >> 1/N_e_), we briefly explore the case where migration is weak relative to drift. As long as the migration rate is sufficiently low (i.e., m << 1/Ne), it is expected that the conditionally neutral mutations will be purged (for CD mutations) or will fix (for CB mutations) in the patch where s ≠ 0, and will operate as neutral mutations in the patch where s ≠ 0. For CD mutations, we would expect them to accumulate in the patch where they are neutral at a rate that is roughly proportional to the mutation rate, as per the classic neutral molecular clock result (Kimura 1968). When we simulated cases with CD mutations, we indeed saw that the rate of accumulation of GxE for fitness was approximately linear (Figure S8), in contrast to the apparent state of equilibrium that was observed at high migration (e.g. Figure S2). The rate of accumulation of non-local load in this case scales with mutation rate, consistent with above prediction, and also increases with increasing strength of selection. It is clear from these simulations that the accumulation CD mutations when drift is strong relative to migration will eventually lead to speciation, when the average fitness difference between home and away is large enough to be lethal.

### Genomic signatures of transient local adaptation and non-local load

Detecting the genomic basis of local adaptation typically involves scanning for loci with elevated allele frequency differentiation among populations (Lewontin and Krakauer 1973; Hoban et al. 2016). To explore whether a genomic signature can be found for transient local adaptation driven by CB alleles and non-local load driven by CD alleles, we calculated F_ST_ for all neutral and fitness-affecting loci in the final generation of the simulations. Using F_ST_ at neutral loci as a null distribution, we found that CB simulations tended show an enrichment of fitness-affecting loci with extreme values of F_ST_ when fitness differences between home and away were most pronounced (Figure 4 A&B). This occurs because, in the patch where they are favoured, the CB mutations tend to increase in frequency (before eventually fixing globally) faster than new neutral mutations, and F_ST_ is elevated while they are transiently segregating. In contrast, CD simulations tended to have relatively fewer fitness-affecting loci falling in the 1% tail of the distribution of F_ST_ for neutral alleles when non-local maladaptation was most pronounced (Figure 4 C&D). While the particular CD alleles responsible for driving most of the home vs. away fitness effect did tend to have the highest F_ST_ values (as this is necessary to cause the fitness difference), these alleles still didn’t usually surpass F_ST_ found for neutral loci (Figure 4 E&F). Thus, it will be very difficult to use conventional genome scan approaches to identify the loci responsible for non-local maladaptation, but potentially much easier for such methods to identify causal CB loci (but see Yoder and Tiffin 2018).

**Figure 4.**
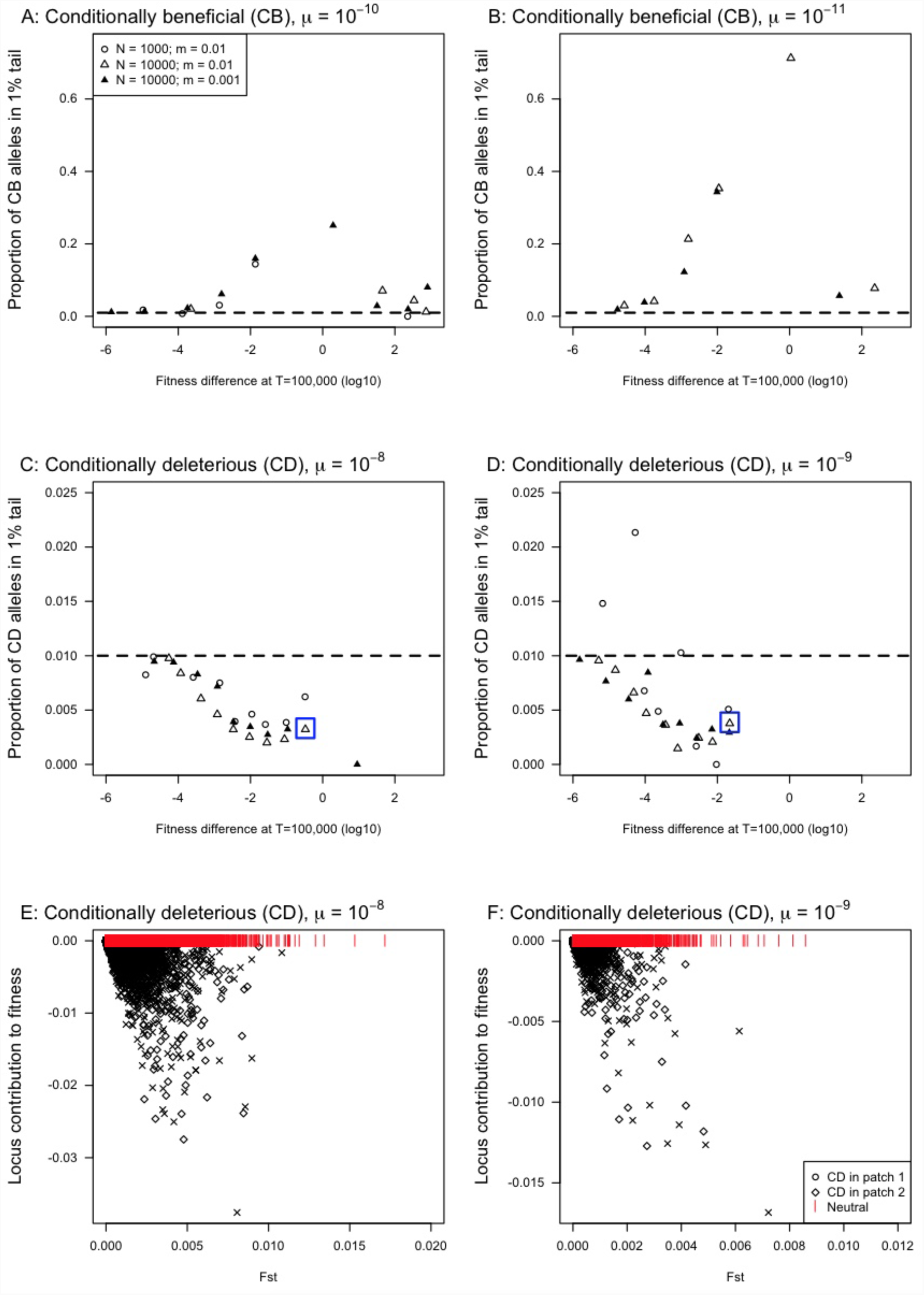
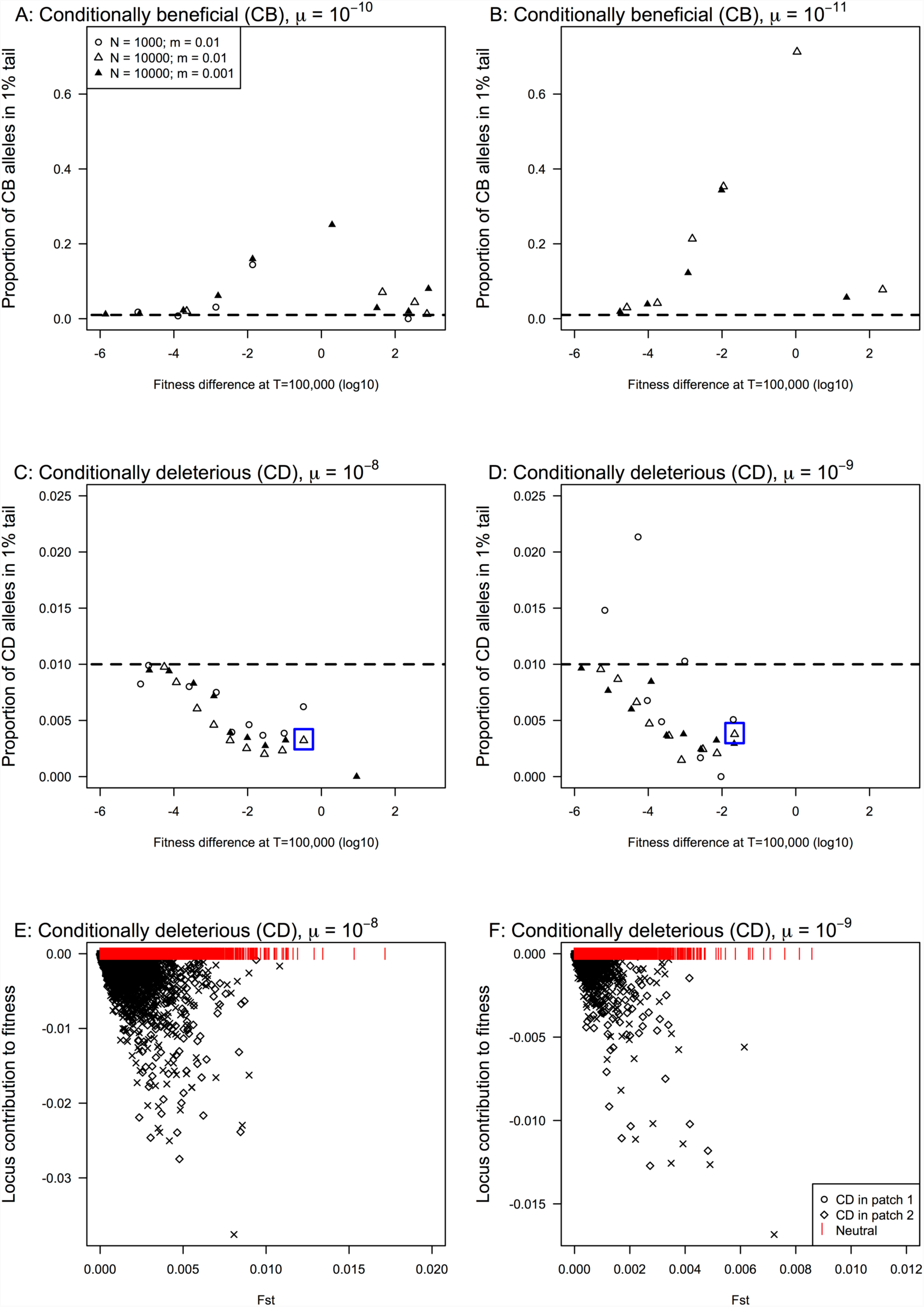
Extreme signatures of population genetic differentiation (F_ST_) are common in simulations with conditionally beneficial mutations (CB), but are rare in simulations with conditionally deleterious mutations (CD). Panels A-D show the proportion of fitness-affecting mutations with FST values that fall into the 1% tail of the distribution of F_ST_ values found for neutral loci, as a function of the average fitness difference for individuals at home vs. away. Each point represents the average across all replicates for simulations run under a given mean selection coefficient; panels A & B show results for Conditionally Beneficial (CB) mutations, while panels C & D show results for Conditionally Deleterious (CD) mutations. Panels E & F illustrate the relationship between the fitness effect of a given locus and its F_ST_ value for the cases in blue squares C & D (N = 10,000 and m = 0.01).

In the CD simulations, the proportion of fitness-affecting loci falling into the 1% tail of the neutral F_ST_ distribution declined below 1% as the strength of selection increased (Figure 4 C&D). We also observed a decline in the ratio of segregating CD to neutral mutations with increasing home-away fitness differences (Figure 5 A&B), and a decline in sojourn time and density with increasing mean selection coefficients (supporting material, Figure S9). Hence, despite the fact that increasing the mean selection coefficient in our simulations resulted in more mutations with strong deleterious effects, the largest mutations (with the strongest effects) were purged very quickly and did not accumulate sufficiently to contribute substantially to home-away fitness differences. So, the relevant effect of a higher mean selection coefficient was an increase in the number of moderately deleterious mutations with sufficiently long sojourn times and densities to have a cumulative effect on home-away fitness differences. The decline in the proportion of fitness-affecting loci falling into the 1% tail of the neutral F_ST_ distribution was, therefore, driven by purging of large-effect loci (which were more abundant when mean *s* was high), while the extent of home-away fitness differences was driven by the accumulation of fitness-affecting loci with moderate fitness effects (which were also more abundant when mean *s* was high). The robustness of these patterns to different assumptions about the distribution of fitness effects is a potential avenue for future research, but we expect that any realistic distribution will have an increased proportion of large-effect mutations when the mean selection coefficient is higher. As long as this basic feature holds, our results should not change qualitatively with different assumptions regarding the distribution of mutation effect size.

**Figure 5.**
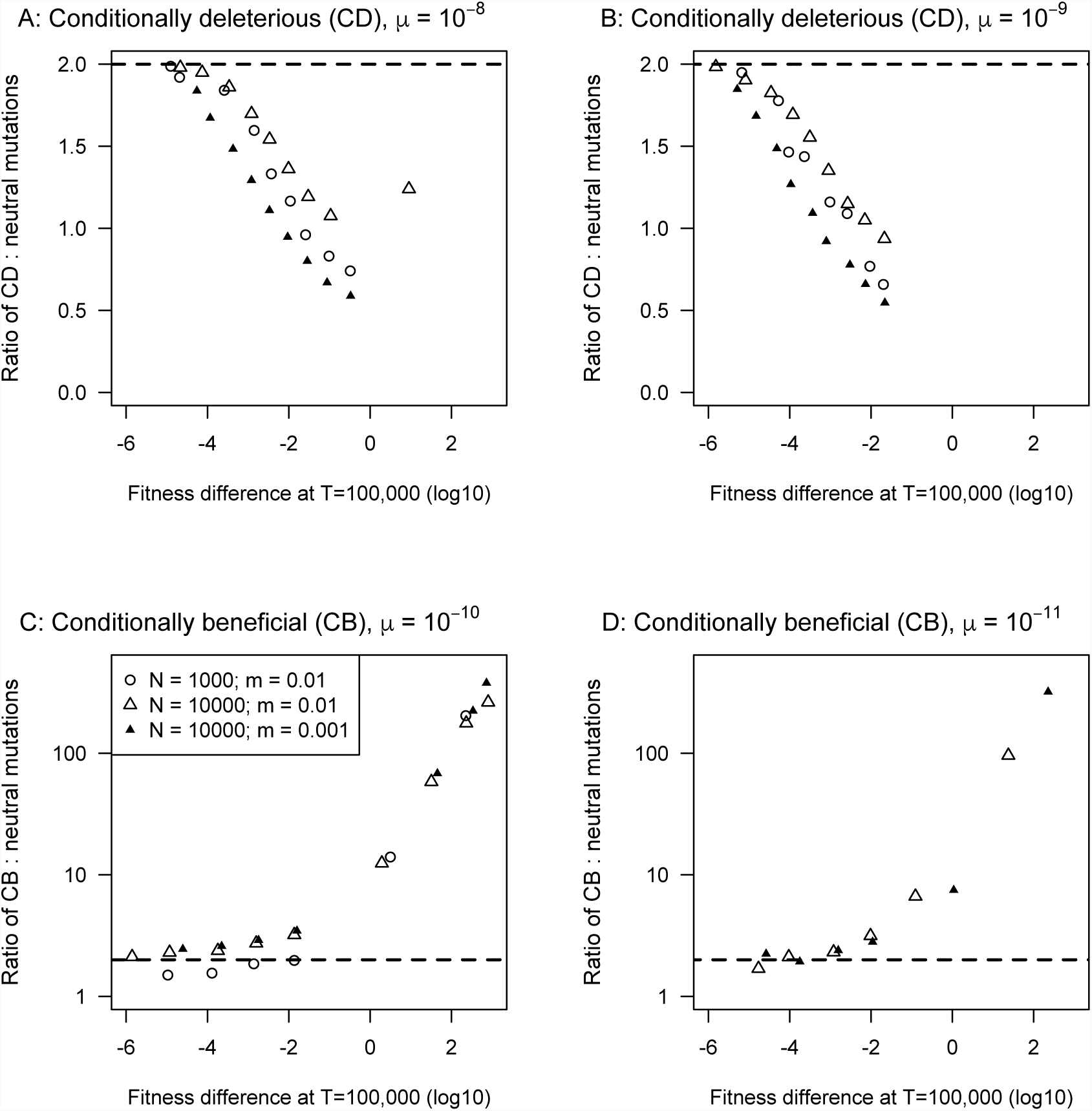
The number of fitness-affecting mutations relative to neutral mutations varies with the fitness difference between home vs. away environments in opposite ways for simulations with Conditionally Deleterious mutations (Panels A & B) and Conditionally Beneficial mutations (Panels C & D). In all simulations, the mutational target for fitness-affecting loci is twice that of neutral loci, indicated by the dashed horizontal line. Each point represents the average across all replicates for simulations run under a given mean selection coefficients.

## Discussion

Local adaptation is most commonly associated with antagonistically pleiotropic mutations (Felsenstein 1976; Savolainen et al. 2013; Bürger 2014). Our study suggests that conditionally neutral mutations, and in particular conditionally deleterious mutations, result in evolutionary dynamics that may play an important role contributing to the genotype by environment interaction for fitness in transplant experiments that is commonly interpreted as evidence of local adaptation (Hedrick et al. 1976; Linhart and Grant 1996; Hereford 2009). The amount of load observed in simulations is well-described by Eq. (2), despite the many simplifying assumptions involved in its derivation, providing an intuitive description of the importance of the various evolutionary processes involved. Thus, a given home-away fitness difference observed in a transplant experiment could, hypothetically, be caused by non-local maladaptation due to the accumulation of non-local load resulting from CD mutations or by transient local adaptation resulting from CB mutations. Unlike the high allele frequency divergence expected in particular cases when antagonistic pleiotropy is involved in local adaptation, which produces strong signatures of selection in genome scans, the loci involved in driving fitness effects could be numerous, transient, and practically undetectable in genome scans when CD mutations are involved in non-local maladaptation. Our results reinforce the conclusion that genome scans have limited power to detect conditionally neutral that contribute to local adaptation (Yoder and Tiffin, 2018).

The amounts of non-local maladaptation observed here are relatively minor in absolute terms (<10% difference in fitness between home and away) and are unlikely to explain a large fraction of observed fitness differences commonly attributed to local adaptation on their own. We have illustrated, nonetheless, that CD mutations may be a factor contributing to apparent signatures of local adaptation that is commonly overlooked in the focus on antagonistic pleiotropy (or beneficial mutations in general). Conditionally deleterious mutations may be a particularly potent factor when combined with AP alleles, since the latter will tend to “protect” flanking regions from gene flow (Barton and Bengtsson 1986; Charlesworth et al. 1997) and will therefore facilitate the accumulation of CD mutations in flanking regions. This effect would be particularly strong for large-effect AP alleles, as selection on such loci results in reduced effective migration rate over a wider region of the chromosome (Barton 2000).

The role of conditionally deleterious mutations in patterns of local adaptation may be quite important because these types of mutations are likely common. Deleterious mutations are much more common than beneficial mutations, and it is reasonable to assume that there would be some condition-dependence in their deleterious effects. For simplicity, we studied conditional mutations that are exactly neural in one environment (as shown in figure 1). In practice, our results would hold if these mutations were nearly-neutral in one environment relatively strongly selected in the other - as long as the selection coefficient in the “neutral” environment is below the selection-drift threshold, these mutations will behave like exactly neutral mutations. So, we contend that, taken together, conditionally nearly-neutral and conditionally exactly-neutral mutations may be common relative to antagonistic pleiotropy mutations with strong and opposite effects in alternative environments.

As an empirical strategy, it has been common for studies of local adaptation to infer conditional neutrality when they fail to observe a fitness effect in one environment of a reciprocal transplant experiment (*e.g.* Bono et al. 2017). However, this should not be interpreted as proof of conditional neutrality (due to either CB or CD) because this could be a consequence of either insufficient power to detect a fitness effect or an experimental design that did not include the causal factor in the natural environment. Common garden experiments are necessarily limited in scope and may exclude ecologically important factors such as density effects, competitors, or rare extreme weather, which could all constitute unsampled events driving fitness differences between genotypes. Furthermore, natural selection can be highly effective even with small fitness effect sizes (*s* acting on alleles on the order of 1/Ne; Crow and Kimura 1970), and it is quite plausible that antagonistic pleiotropy could be operating despite being nearly impossible to detect with sample sizes of only a few tens or hundreds of individuals. Because of the nature of scientific evidence (*i.e*. it is impossible to conclusively prove a negative), it is practically impossible to conclusively show that some form of conditional neutrality (either CB or CD) is operating in nature, as this requires demonstrating that there is no fitness difference between genotypes in one location, and it is always possible that some important causal factor has not been included in the experimental design. Even if we could reliably detect conditionally dependent selection (CD or CB) using transplant experiments, it would still be very difficult to differentiate CB from CD given a snapshot of fitness differences or mutation effect sizes and directions. Combining outlier scans with orthogonal lines of evidence, such as predictive approaches to identify deleterious mutations (*e.g*. Ng and Henikoff 2001; Choi et al. 2012), may increase the power to discriminate conditionally deleterious vs. neutral alleles falling in the tails of the distribution. Finally, because deleterious mutations are an important driver of disease in humans, understanding how such mutations accumulate in different regions of the genome may have important applications in medicine.

## Acknowledgements

We would like to thank J. Travis, M. Williamson, D. Lindtke, K. Hodgins, and two anonymous reviewers for helpful comments and suggestions during the preparation of this manuscript. This work was funded by an AIHS grant and enabled in part by computational support provided by Compute Canada.

## Supplementary Figures

**Figure S1:**
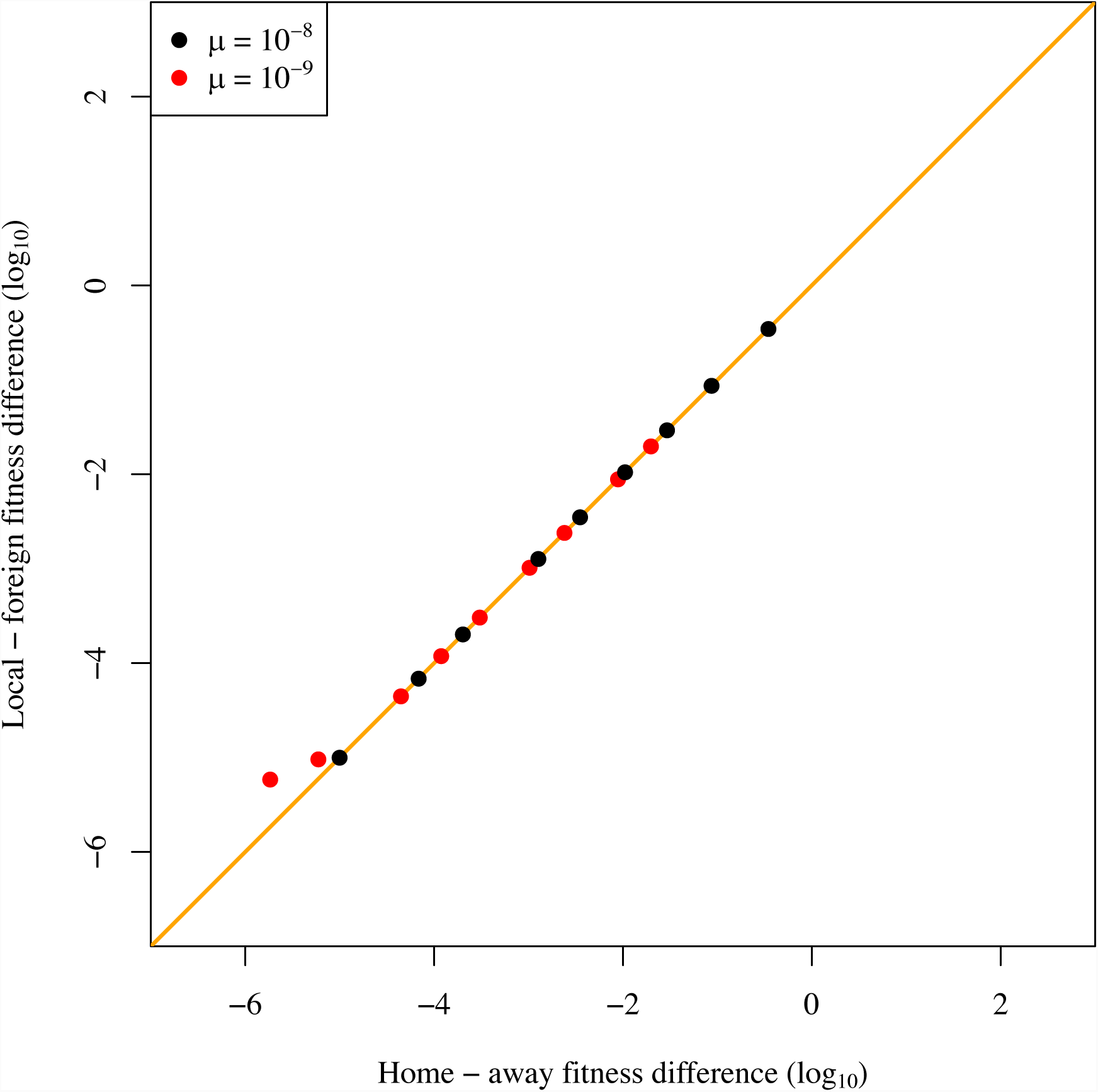
Comparison of home versus away fitness differences and local versus foreign fitness differences. The orange line indicates the 1:1 relationship. For this plot, m = 0.01 and N = 1000.

**Figure S2:**
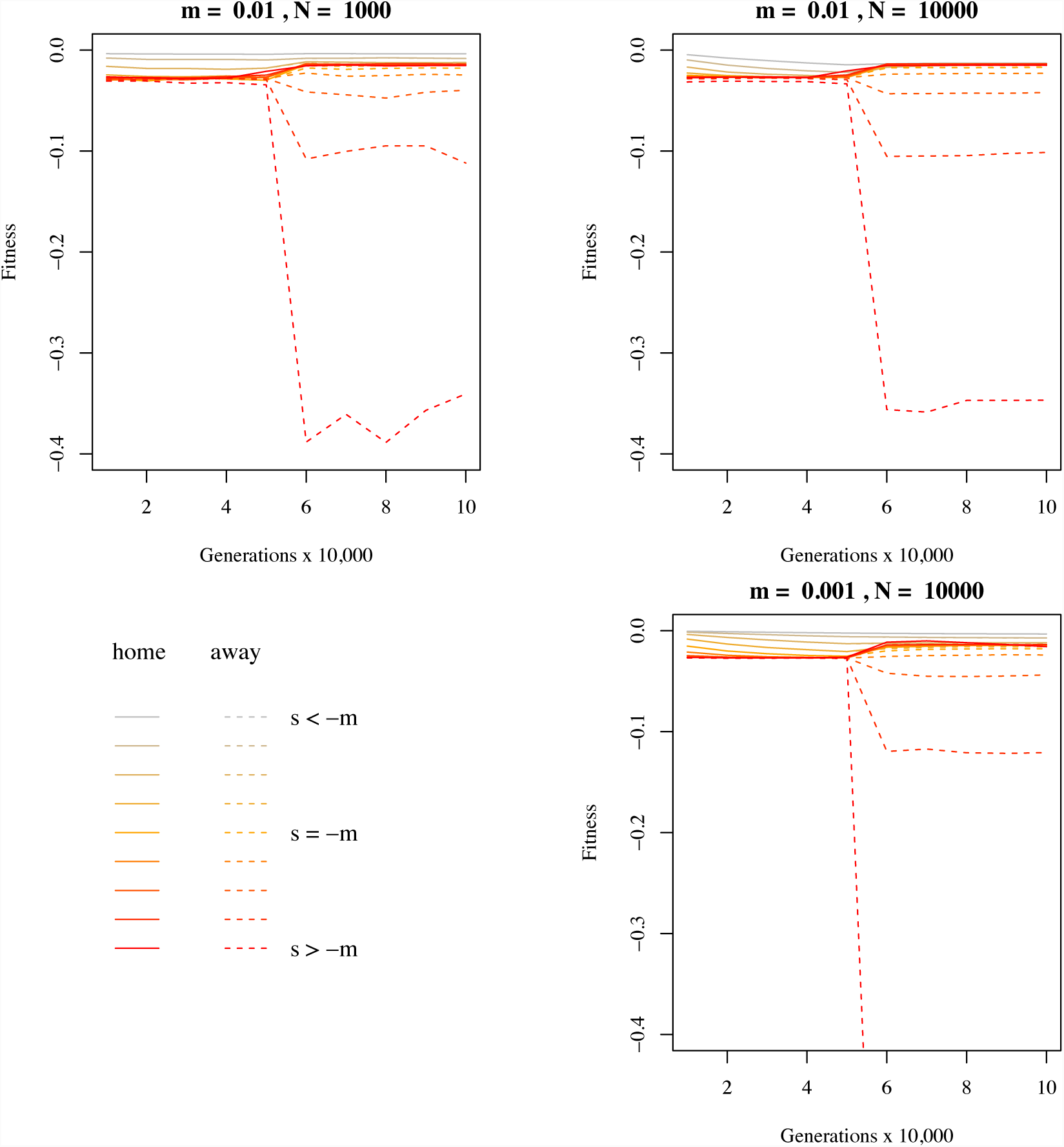
The average home-away fitness difference (averaged across replicates) at each sampled time-point in simulations with conditionally deleterious mutations and μ = 10^−8^. The difference between home fitness (solid lines) and away fitness (dashed lines), averaged over the last 40,000 generations, was used in subsequent analyses.

**Figure S3:**
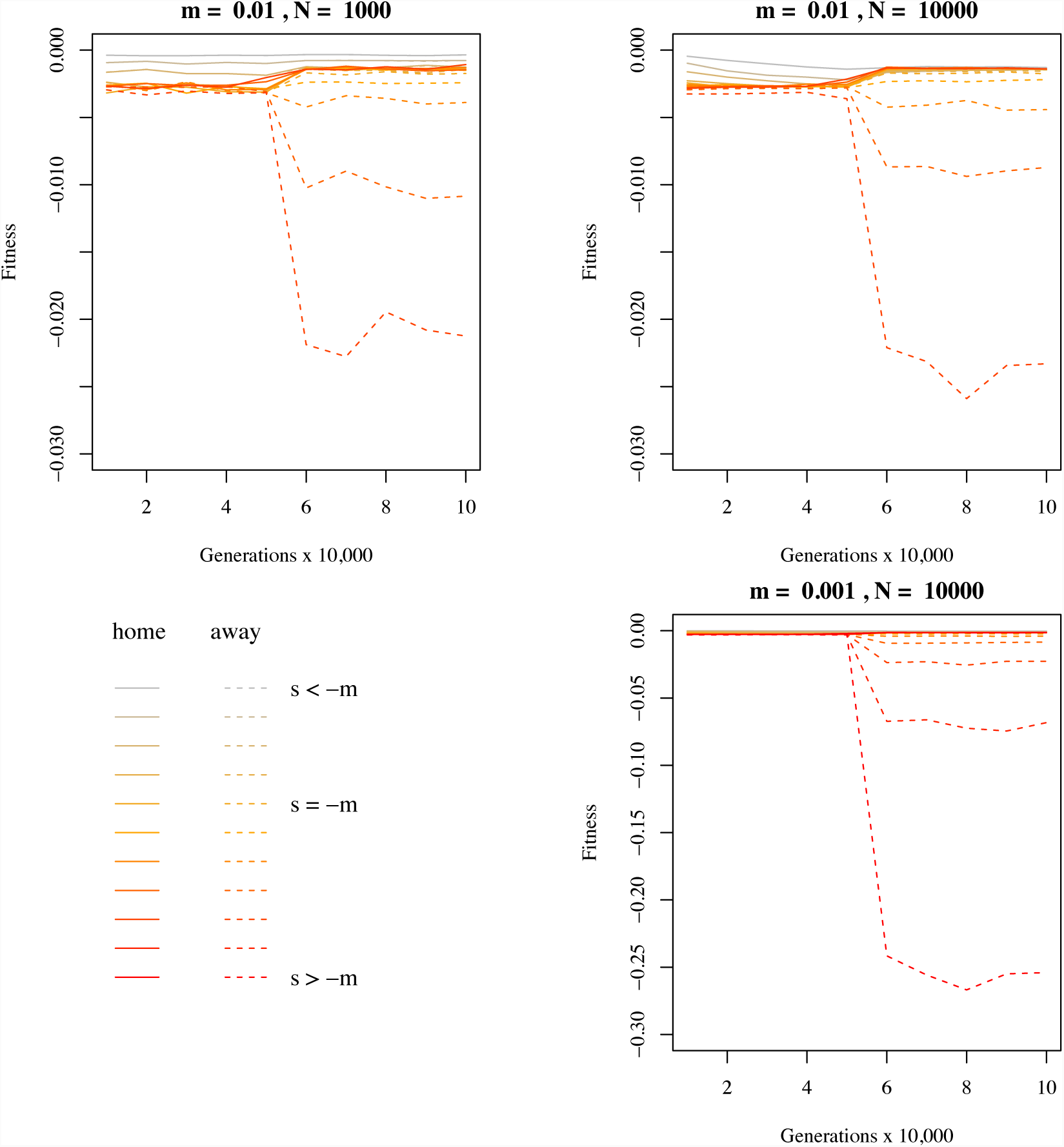
The average home-away fitness difference (averaged across replicates) at each sampled time-point in simulations with conditionally deleterious mutations and μ = 10^−9^. The difference between home fitness (solid lines) and away fitness (dashed lines), averaged over the last 40,000 generations, was used in subsequent analyses.

**Figure S4:**
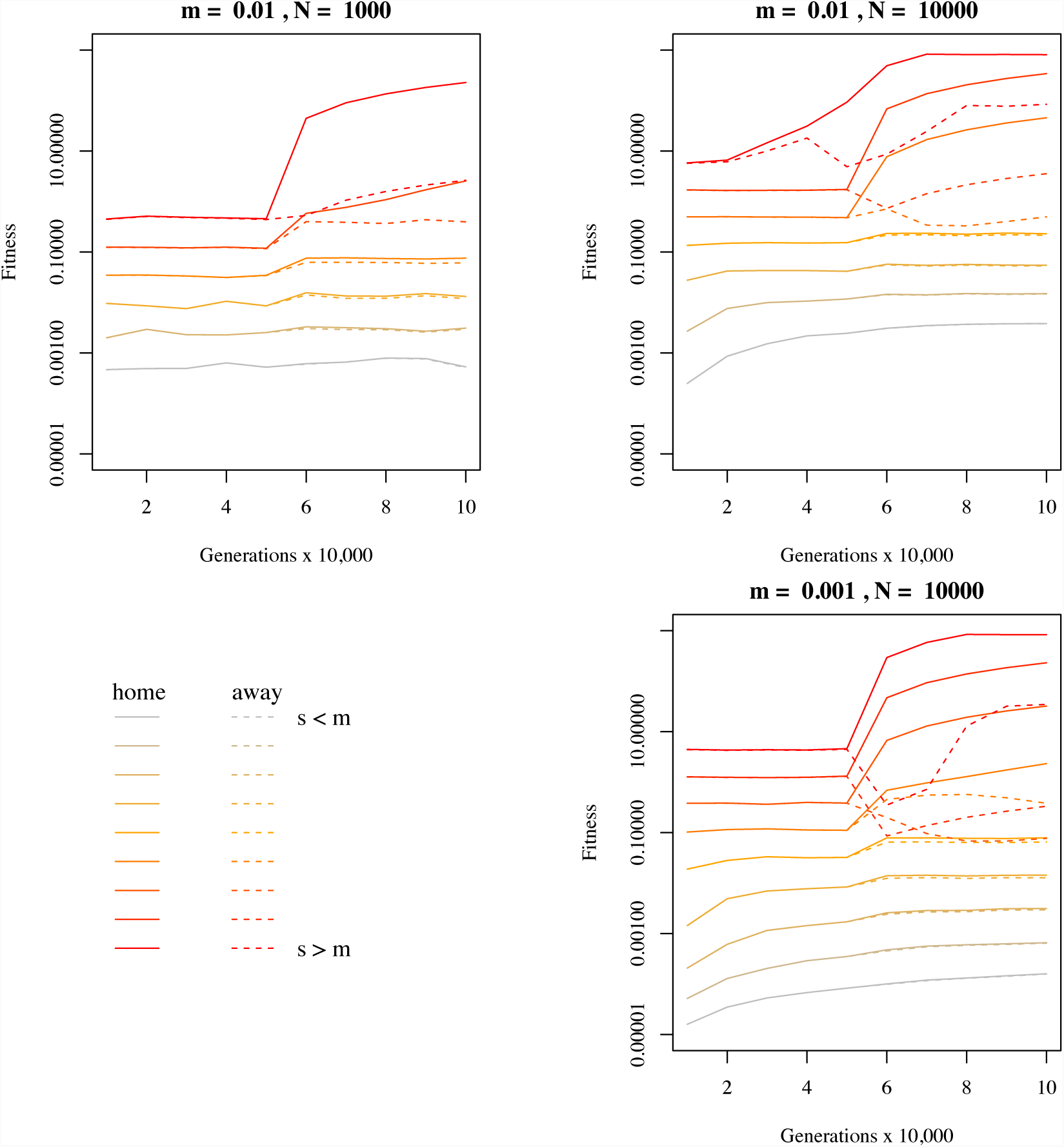
The average home-away fitness difference (averaged across replicates) at each sampled time-point in simulations with conditionally beneficial mutations and μ = 10^−10^. The difference between home fitness (solid lines) and away fitness (dashed lines), averaged over the last 40,000 generations, was used in subsequent analyses.

**Figure S5:**
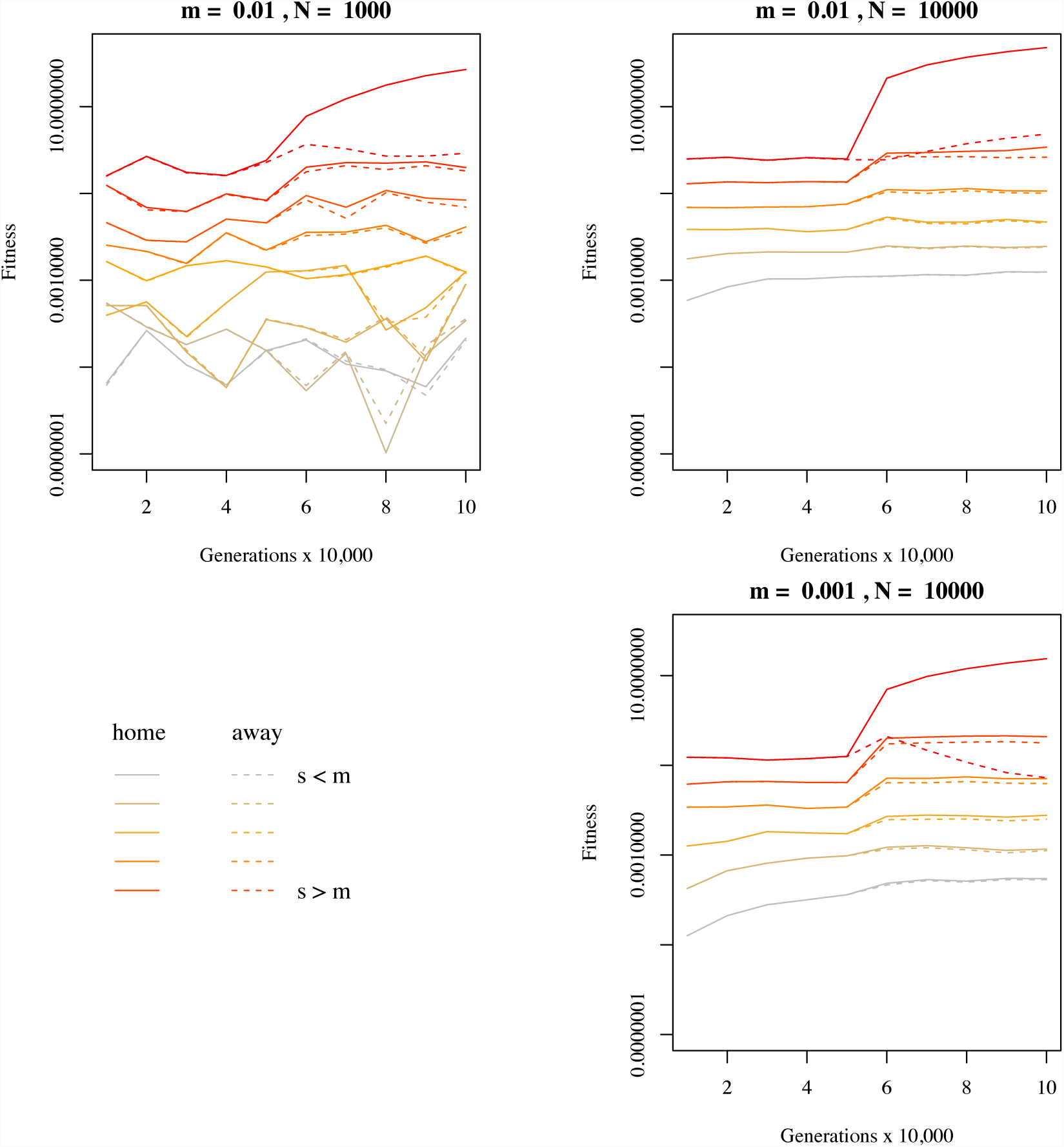
The average home-away fitness difference (averaged across replicates) at each sampled time-point in simulations with conditionally beneficial mutations and μ = 10^−11^. The difference between home fitness (solid lines) and away fitness (dashed lines), averaged over the last 40,000 generations, was used in subsequent analyses.

**Figure S6:**
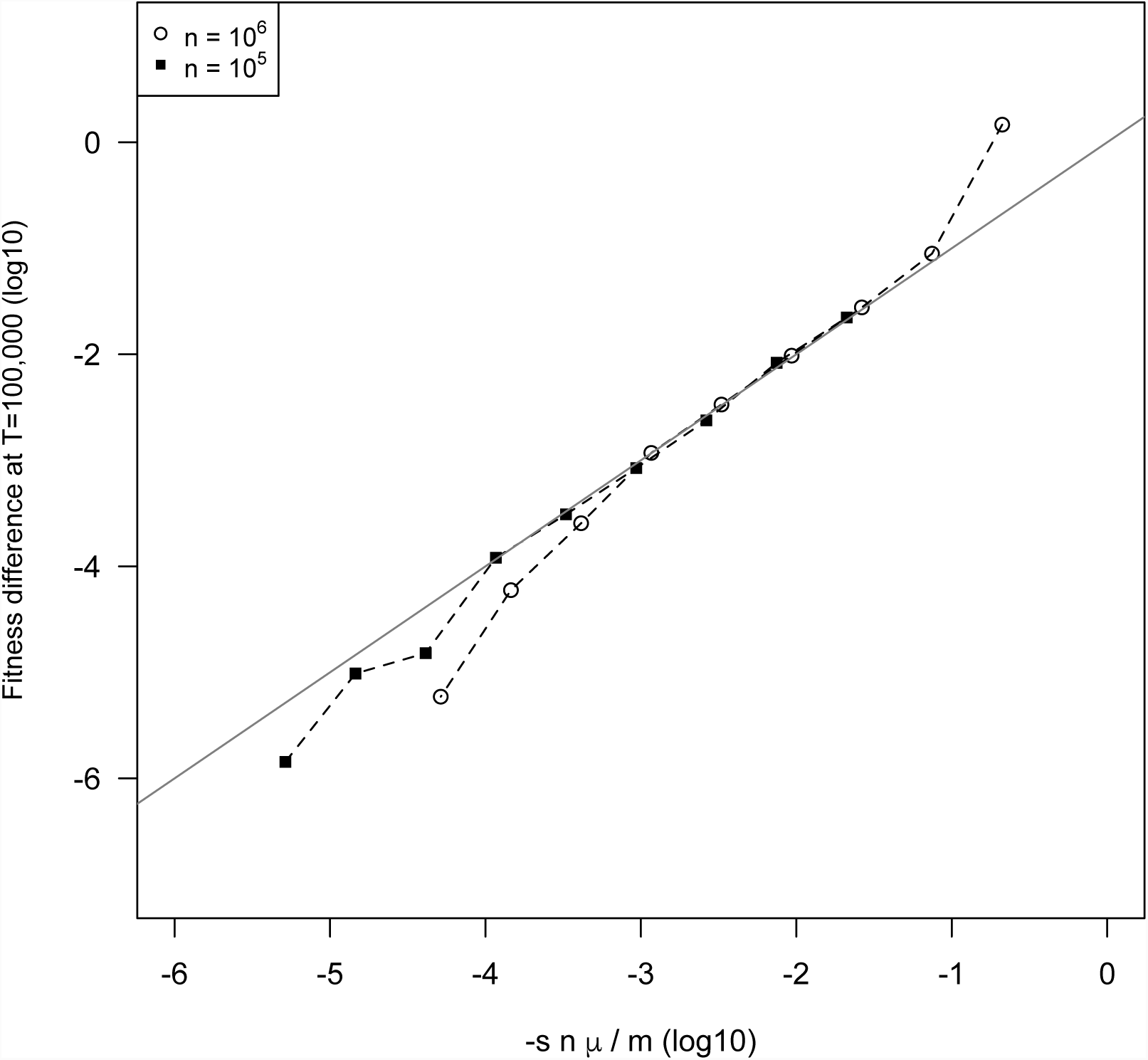
Reducing the number of mutable sites by a factor of 10 does not substantially affect the linear relationship between the prediction of equation (1) and the simulation results, except at very low predicted fitness differences. Simulations have *N* = 1000, *m* = 0.01, *r* = 10^−6^, and μ = 10^−8^ and either *n* = 10^6^ or *n* = 10^5^ mutable sites.

**Figure S7:**
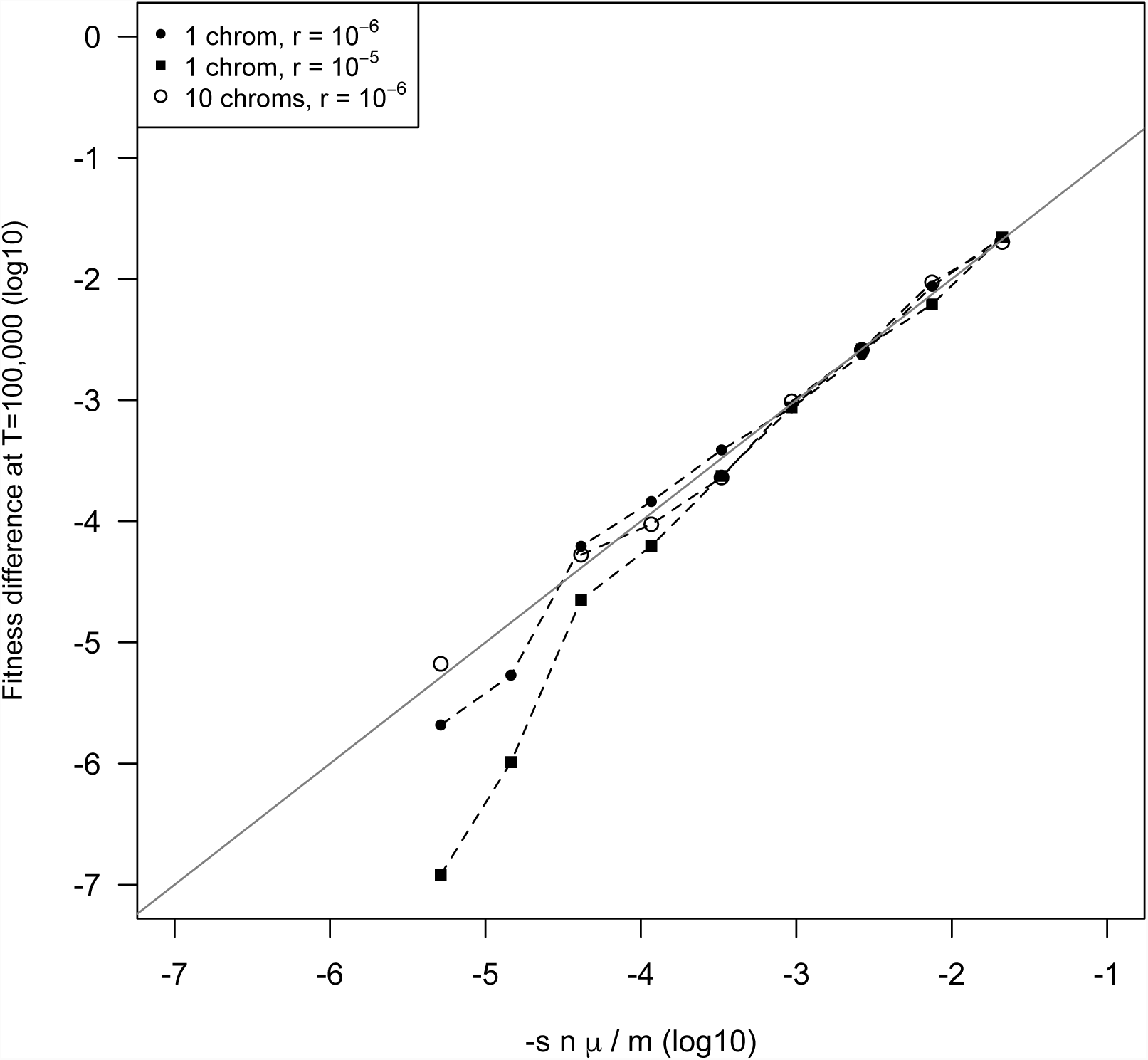
Changing the number of chromosomes and recombination rate among adjacent loci does not substantially affect the linear relationship between the prediction of equation (1) and the simulation results, except at very low predicted fitness differences. Simulations have *N* = 1000, *m* = 0.01, *n* = 10^6^ sites, and μ = 10^−9^, and either 1 chromosome with recombination rate of *r* = 10^−5^ or 10^−6^, or 10 chromosomes with *r* = 10^−6^ between adjacent sites.

**Figure S8:**
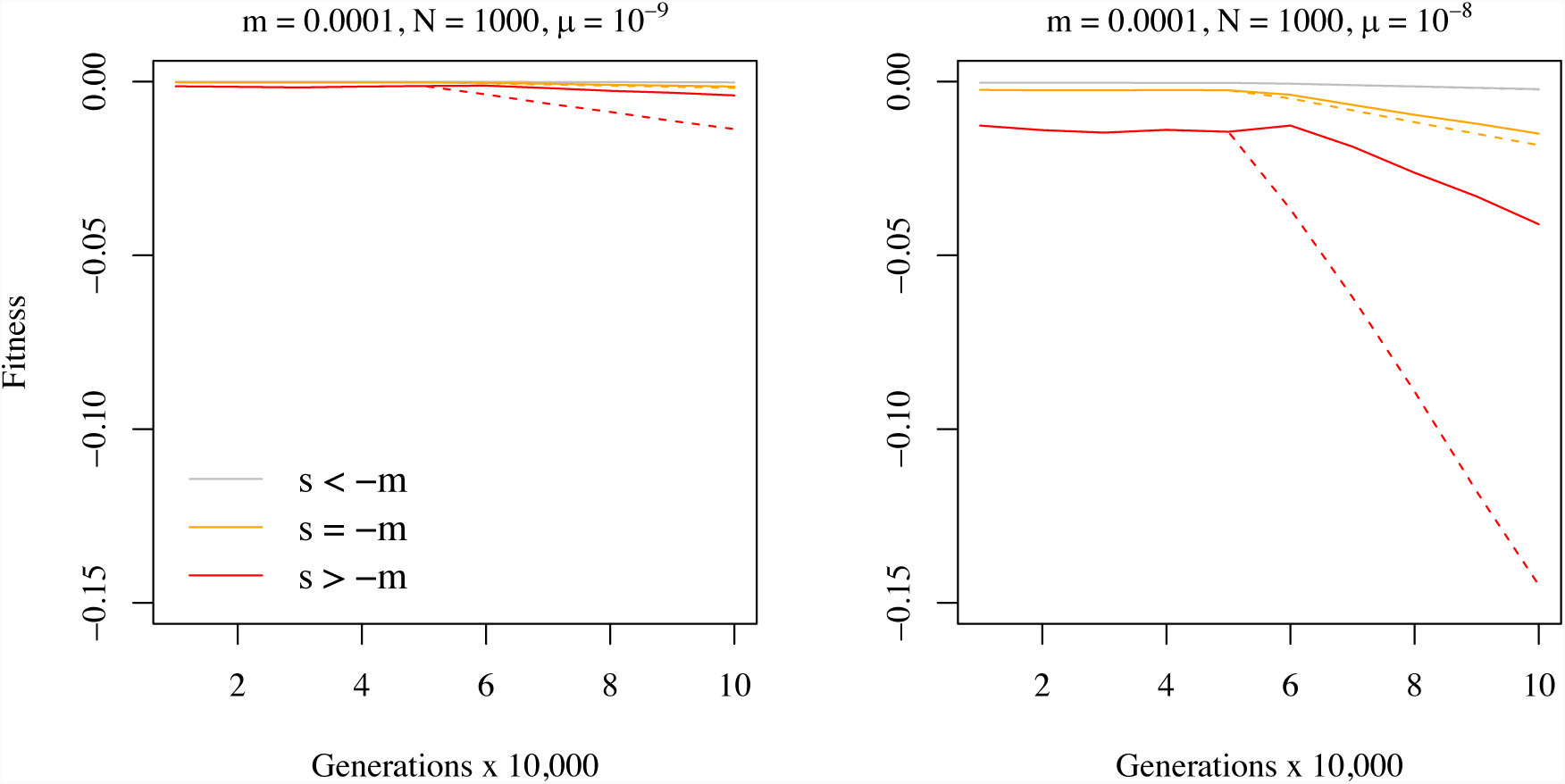
Accumulation of CD mutations causing differences in fitness between home (solid) and away (dashed) environments, when migration is weak relative to drift (m = *N*/10) for different mutation rates and mean strengths of selection.

**Figure S9:**
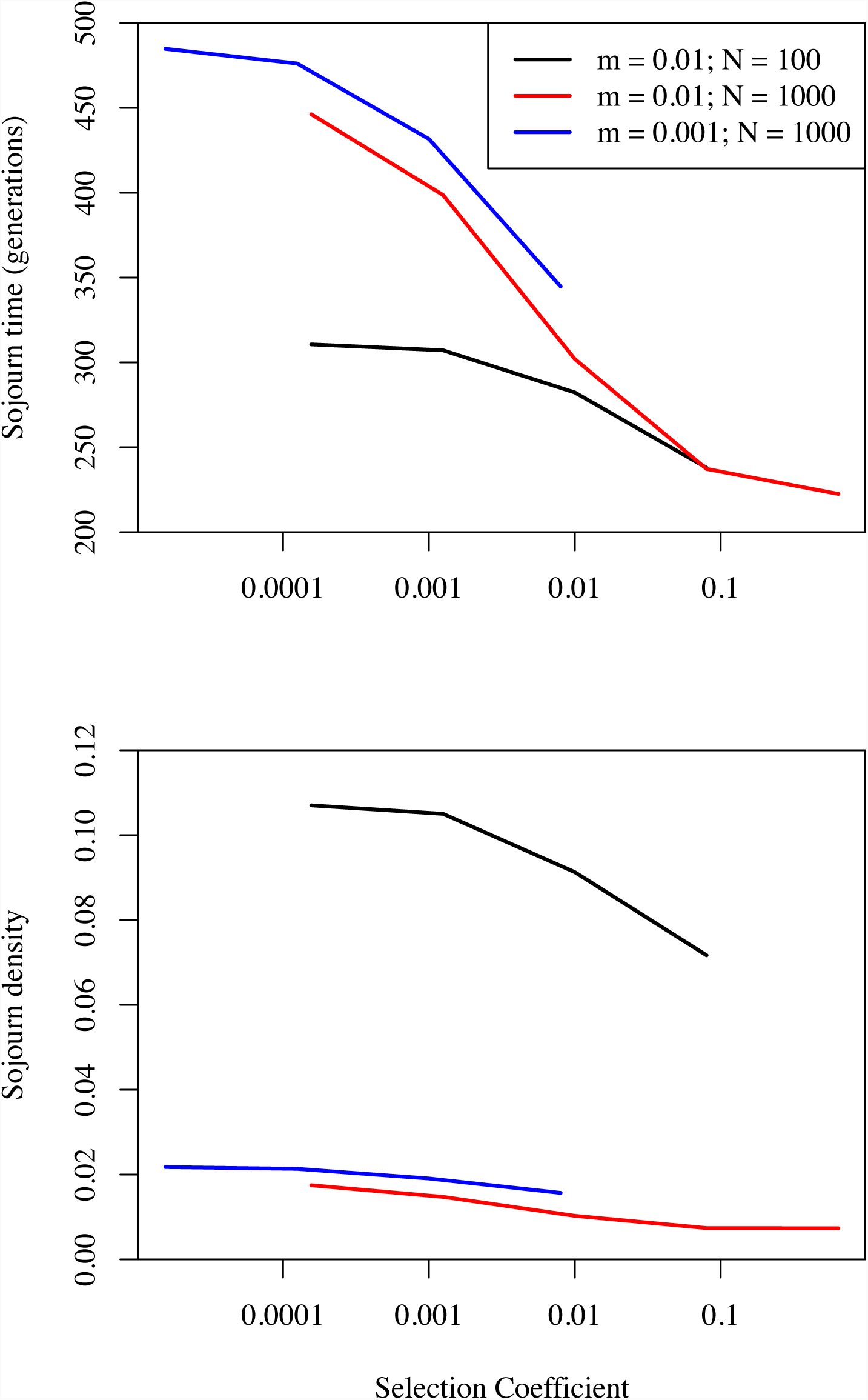
The sojourn time and sojourn density for conditionally deleterious mutations in simulations with μ = 10^−8^.

## Literature cited

Agrawal, A.F., and M.C. Whitlock. 2012. Mutation load: the fitness of individuals in populations where deleterious alleles are abundant. Annual Review of Ecology, Evolution, and Systematics 43: 115–135.

Anderson, J. T., C.-R. Lee, C. a Rushworth, R. I. Colautti, and T. Mitchell-Olds. 2013. Genetic trade-offs and conditional neutrality contribute to local adaptation. Molecular ecology 22: 699–708.

Barton, N. 2000. Genetic hitchhiking. Philosophical transactions of the Royal Society of London. Series B, Biological sciences 355: 1553–1562.

Barton, N., and B. O. Bengtsson. 1986. The barrier to genetic exchange between hybridising populations. Heredity 57: 357–376.

Bataillon, T. 2000. Estimation of spontaneous genome-wide mutation rate parameters: whither beneficial mutations? Heredity 84: 497–501.

Blanquart, F., O. Kaltz, S. L. Nuismer, and S. Gandon. 2013. A practical guide to measuring local adaptation. Ecology Letters 16: 1195–1205.

Bono, L. M., Smith, L. B., Pfennig, D. W. and Burch, C. L. (2017). The emergence of performance trade-offs during local adaptation: insights from experimental evolution. Molecular Ecology 26: 1720–1733.

Bulmer, M. 1980. The mathematical theory of quantitative genetics. Clarendon Press, Oxford, UK.

Bürger, R. 2000. The mathematical theory of selection, recombination, and mutation. John Wiley & Sons, Chichester, UK.

Bürger, R. 2014. A survey of migration-selection models in population genetics. Discrete and Continuous Dynamical Systems - Series B 19 (4): 883–959.

Charlesworth, B., M. Nordborg, and D. Charlesworth. 1997. The effects of local selection, balanced polymorphism and background selection on equilibrium patterns of genetic diversity in subdivided populations. Genetical research 70: 155–174.

Choi, Y., G. E. Sims, S. Murphy, J. R. Miller, and A. P. Chan. 2012. Predicting the Functional Effect of Amino Acid Substitutions and Indels. PLoS ONE 7.

Crow, J.F. and Kimura, M. 1970. An introduction to population genetic theory. Harper and Row, New York.

Felsenstein, J. 1976. The theoretical population genetics of variable selection and migration. Annual Review of Genetics. 10: 253–280.

Fournier-Level, A., A. Korte, M. D. Cooper, M. Nordborg, J. Schmitt, and A. M. Wilczek. 2011. A Map of Local Adaptation in Arabidopsis thaliana. Science 334: 86–89.

Fry, J.D. 1992. On the maintenance of genetic variation by disruptive selection among hosts in a phytophagous mite. Evolution 46: 279–283.

Fry, J. D. 1996. The University of Chicago The Evolution of Host Specialization: Are Trade-Offs Overrated? Most species of phytophagous insects The reasons evolves and is maintained leads to the prediction species should be specialized on the same hosts. Even American. The American naturalist 148: S84–S107.

Haldane, J.B.S. 1927. A Mathematical Theory of Natural and Artificial Selection, Part V: Selection and Mutation. Mathematical Proceedings of the Cambridge Philosophical Society 23 (07): 838–844.

Hall, M. C., D. B. Lowry, and J. H. Willis. 2010. Is local adaptation in Mimulus guttatus caused by trade-offs at individual loci? Molecular Ecology 19: 2739–2753.

Hedrick, P., Ginevan, M.E., Ewing, E.P. 1976. Genetic polymorphism in heterogeneous environments. Annual Review of Ecology and Systematics. 7: 1–32.

Hereford, J. 2009. A Quantitative Survey of Local Adaptation and Fitness Trade-Offs. American Naturalist 173: 579–588.

Hoban, S., J. L. Kelley, K. E. Lotterhos, M. F. Antolin, G. Bradburd, D. B. Lowry, M. L. Poss, et al. 2016. Finding the Genomic Basis of Local Adaptation: Pitfalls, Practical Solutions, and Future Directions. The American Naturalist 188: 379–397.

Kawecki, T. J. 1997. Sympatric Speciation via Habitat Specialization Driven by Deleterious Mutations. Evolution 51: 1751–1763.

Kawecki, T. J., and D. Ebert. 2004. Conceptual issues in local adaptation. Ecology Letters 7: 1225–1241.

Keightley, P. D., and A. Eyre-Walker. 2010. What can we learn about the distribution of fitness effects of new mutations from DNA sequence data? Philosophical Transactions of the Royal Society B: Biological Sciences 365: 1187–1193.

Kimura, M. 1968. Evolutionary rate at the molecular level. Nature 217 (5129): 624–626.

Lewontin, R. C., and J. Krakauer. 1973. Distribution of gene frequency as a test of the theory of the selective neutrality of polymorphisms. Genetics 74: 175–195.

Linhart, Y.B. and Grant, M.C. Evolutionary significance of local genetic differentiation in plants. Annual Review of Ecology and Systematics. 27: 237–277.

Messer, P. W. 2013. SLiM: Simulating Evolution with Selection and Linkage. Genetics 194: 1037–1039.

Ng, P. C., and S. Henikoff. 2001. Predicting Deleterious Amino Acid Substitutions Predicting Deleterious Amino Acid Substitutions. Genome Research 11: 863–874.

Oakley, C. G., J. Ågren, R. A. Atchison, and D. W. Schemske. 2014. QTL mapping of freezing tolerance: Links to fitness and adaptive trade-offs. Molecular Ecology 23: 4304–4315.

Richardson, J. L., M. C. Urban, D. I. Bolnick, and D. K. Skelly. 2014. Microgeographic adaptation and the spatial scale of evolution. Trends in Ecology and Evolution 29: 165–176.

Savolainen, O., M. Lascoux, and J. Merilä. 2013. Ecological genomics of local adaptation. Nature Reviews Genetics 14: 807–820.

Song, B.H., and Mitchell-Olds, T. 2011. Evolutionary and ecological genomics of non-model plants. Journal of Systematics and Evolution 49: 17–24.

Whitlock, M.C., and Gomulkiewicz, R. 2005. Probability of Fixation in a Heterogeneous Environment Genetics 171: 1407–1417.

Yoder, J.B., and Tiffin, P. 2018. Effects of Gene Action, Marker Density, and Timing of Selection on the Performance of Landscape Genomic Scans of Local Adaptation, Journal of Heredity 109 (1): 16–28. https://doi.org/10.1093/jhered/esx042

